# Comparison of Culture Results between Specimens from Corneal Scraping with Microhomogenization and Corneal Swab in Moderate and Severe Bacterial Corneal Ulcers

**DOI:** 10.1101/2022.09.01.506297

**Authors:** Faraby Martha, Lukman Edwar, Anis Karuniawati, Ahmad Fuady, Ramadhiana Maktazula Tuasikal

## Abstract

**Purpose:** To compare bacterial culture results from corneal infiltrate specimens taken by corneal scrapping followed by microhomogenization and direct scraping methods.

**Methods:** This study was a comparative cross-sectional, with 18 subjects divided into two randomized groups; Group A (first scrapings then swab), and group B (first swab then scrapings). Both scraping and swab methods are separated and then compared in data analysis.

**Results:** The proportion of Gram according to culture in the scraping-microhomogenization technique was 6/13 (46.2%) and the proportion of Gram according to cultures in the swab technique was 5/13 (38.5%). McNemar test p-value is 1.000 (p>0.05). The proportion of positive cultures in the scraping-microhomogenization technique was 13/18 (72.2%) and the proportion of positive culture in swab technique was 9/18 (50%). McNemar test p-value is 0.219 (p> 0.05). The Kappa suitability test value from scraping-microhomogenization technique against the corneal swab is 0.333.

**Conclusion:** The results of bacterial culture from specimens taken by the corneal scraping followed by microhomogenization have a higher positive culture compared to those taken by corneal swab, but not statistically significant.

## Background

Bacterial corneal ulcer is a defect in the corneal epithelium with loss of stroma tissue and/or inflammation of the stroma caused by bacteria.^1,2^ Bacterial infection is the most common cause of corneal ulcers and can cause visual impairment at various ages. The number is expanding with the increasing use of contact lenses, topical steroids, trauma, and abnormalities of the eye surface that cause blindness due to corneal scar, perforation, or endophthalmitis.^3,4^

Examination of bacterial culture as the gold standard treatment for bacterial corneal ulcers is not 100% able to grow germs. It ranges from 37% using a sterile 23 gauge needle and 84% with a flame-sterilized platinum spatula or a calcium alginate swab.^5,6^ In our hospital, positive culture from 220 presumed bacterial corneal ulcer patients was 41.8%. A common method used in the removal of the corneal specimen was a corneal biopsy using a 26 gauge needle in combination with a corneal.^7^

There are two commonly used techniques for taking corneal ulcer specimens, corneal swab and corneal biopsy or scraping.^8^ The corneal swab is more practical and effective in collecting specimens and can provide better bacterial absorption and bacterial release to the solid agar media than spatula does.^8^ Corneal scrapings can take specimens from the corneal edge and the ulcer base, so it is expected that a sufficient amount of bacteria samples is collected and access to infection in deeper corneal stroma is obtained rather than just swabbing the corneal surface.^9^ Corneal scrapings can be combined with the microhomogenization technique. The number of positive cultures is 20% greater in scrapings techniques followed by microhomogenization rather than those using usual technique of corneal scrapings.^10^

Knowing the microorganism that causes corneal infection is very important in the success of therapy. This study was aimed to compare the culture results between scraping specimens with microhomogenization and corneal smears in patients with moderate to severe bacterial corneal ulcers. This study could be a guideline in treating bacterial corneal ulcers.

## Method

This study was a comparative cross-sectional, with 18 subjects divided into two groups after corneal scraping, Group A (scraping-microhomogenization technique), and group B (swab technique). Samples were selected consecutively, with the order in which the samples are randomized.

The inclusion criteria were bacterial corneal ulcer patients (diagnosis based on clinical course) with moderate and severe according to modified Jones classification;^11^ age over 18 years old; willing to participate in the study; and direct smear examinations of Gram and KOH result: bacterial cells were found without fungal hyphae, or no bacterial cells or fungal hyphae were found. The exclusion criteria were patients with clinical perforation or impending perforated bacterial corneal ulcers; endophthalmitis; corneal ulcer which clinically and Gram staining support bacterial corneal ulcers but fungal culture also grow; and samples that cannot be examined in the laboratory for any reason.

The patient was given 0.5% pantocaine eye drops for corneal anesthesia. Then a corneal swab was performed for direct smears for Gram and KOH stains. Corneal scrapings were taken from the edge and base of the ulcer under aseptic conditions using the tip of a 1cc tuberculin syringe bent at 60 degrees under slit lamp magnification and inserted into a sterile container. The corneal swab was taken with a sterile cotton swab and inserted into the Stuart transport medium.

In group A, corneal scraping specimens will be followed by a microhomogenization process until a volume of 80 microliters was obtained and then inoculated on the culture media. In contrast, in group B, corneal swab specimens were directly inoculated onto the culture medium. Both specimens from 2 types of techniques were inoculated on the surface of solid media. The culture medium was incubated at 35°C for 20 hours. The grown isolates were identified biochemically and continued antibiotic susceptibility testing using the Vitek2 automation machine.

## Result

A total of 18 patients were included in this study which divided 9 patients into each group. The number of male subjects was more dominant than those of female subjects but no significant difference between the two groups. There were no significant differences in baseline characteristics of the study subjects between the two groups (Table 1).

**Table 1.**
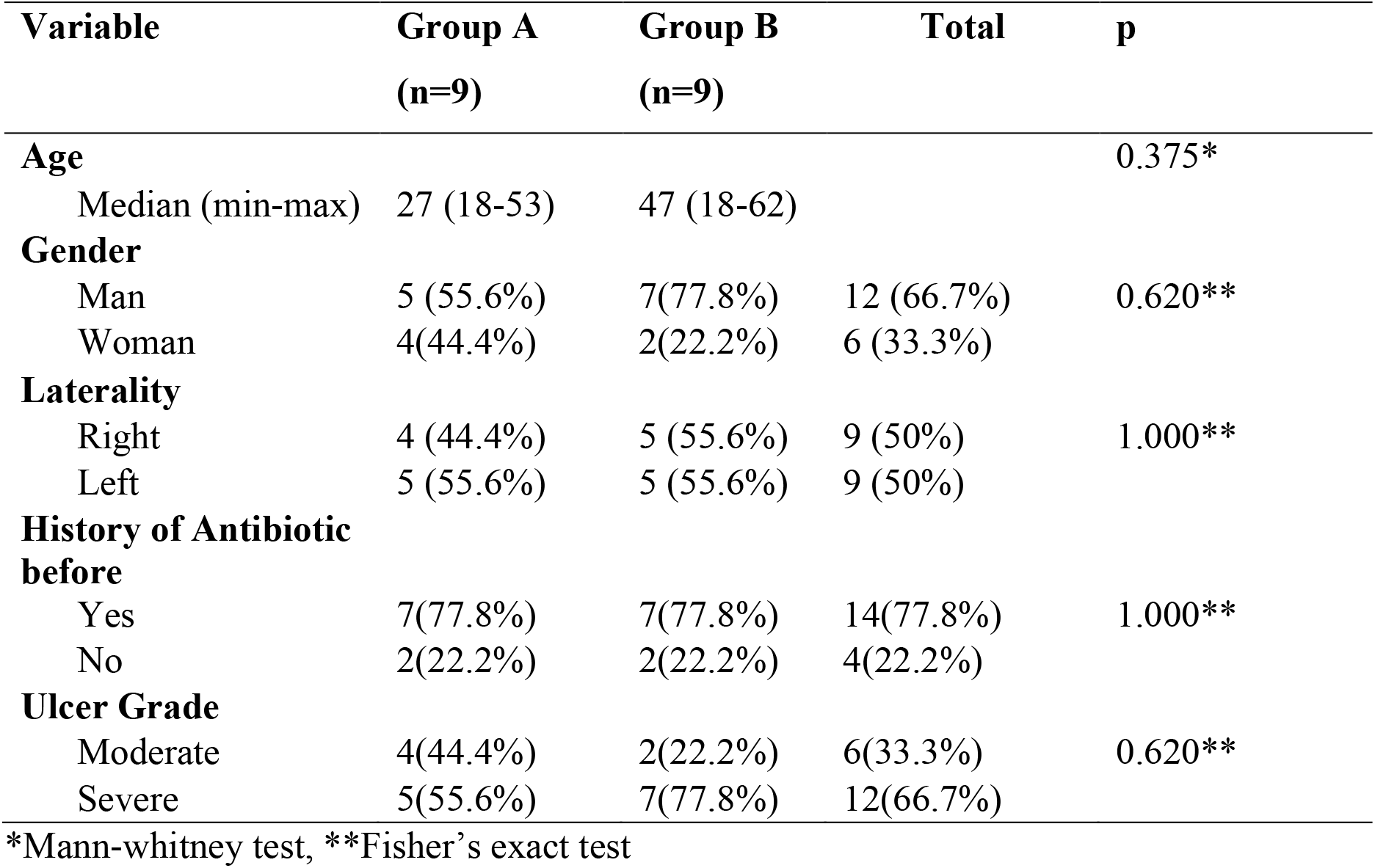
Subject Characteristic (N=18 subject)

| **Variable** | **Group A**<br>**(n=9)** | **Group B**<br>**(n=9)** | **Total** | **p** |
| --- | --- | --- | --- | --- |
| **Age** |  |  |  | 0.375* |
| Median (min-max) | 27 (18-53) | 47 (18-62) |  |  |
| **Gender** |  |  |  | 0.620** |
| Man | 5 (55.6%) | 7(77.8%) | 12 (66.7%) |  |
| Woman | 4(44.4%) | 2(22.2%) | 6 (33.3%) |  |
| **Laterality** |  |  |  | 1.000** |
| Right | 4 (44.4%) | 5 (55.6%) | 9 (50%) |  |
| Left | 5 (55.6%) | 5 (55.6%) | 9 (50%) |  |
| **History of Antibiotic before** |  |  |  | 1.000** |
| Yes | 7(77.8%) | 7(77.8%) | 14(77.8%) |  |
| No | 2(22.2%) | 2(22.2%) | 4(22.2%) |  |
| **Ulcer Grade** |  |  |  | 0.620** |
| Moderate | 4(44.4%) | 2(22.2%) | 6(33.3%) |  |
| Severe | 5(55.6%) | 7(77.8%) | 12(66.7%) |  |

\*Mann-whitney test, \*\*Fisher's exact test

The percentage of positive cultures aims to assess the effectiveness of both examination techniques. Table 2 shows that that the positive culture results of the scraping-microhomogenization technique were higher than swab technique (72.2% vs 50%), which in subject B8 grew 2 bacterias from group A, while no bacteria grew in group B. There were 4 culture results with the same bacteria between the two groups, and two of them were *Pseudomonas aeruginosa*.

**Table 2.**
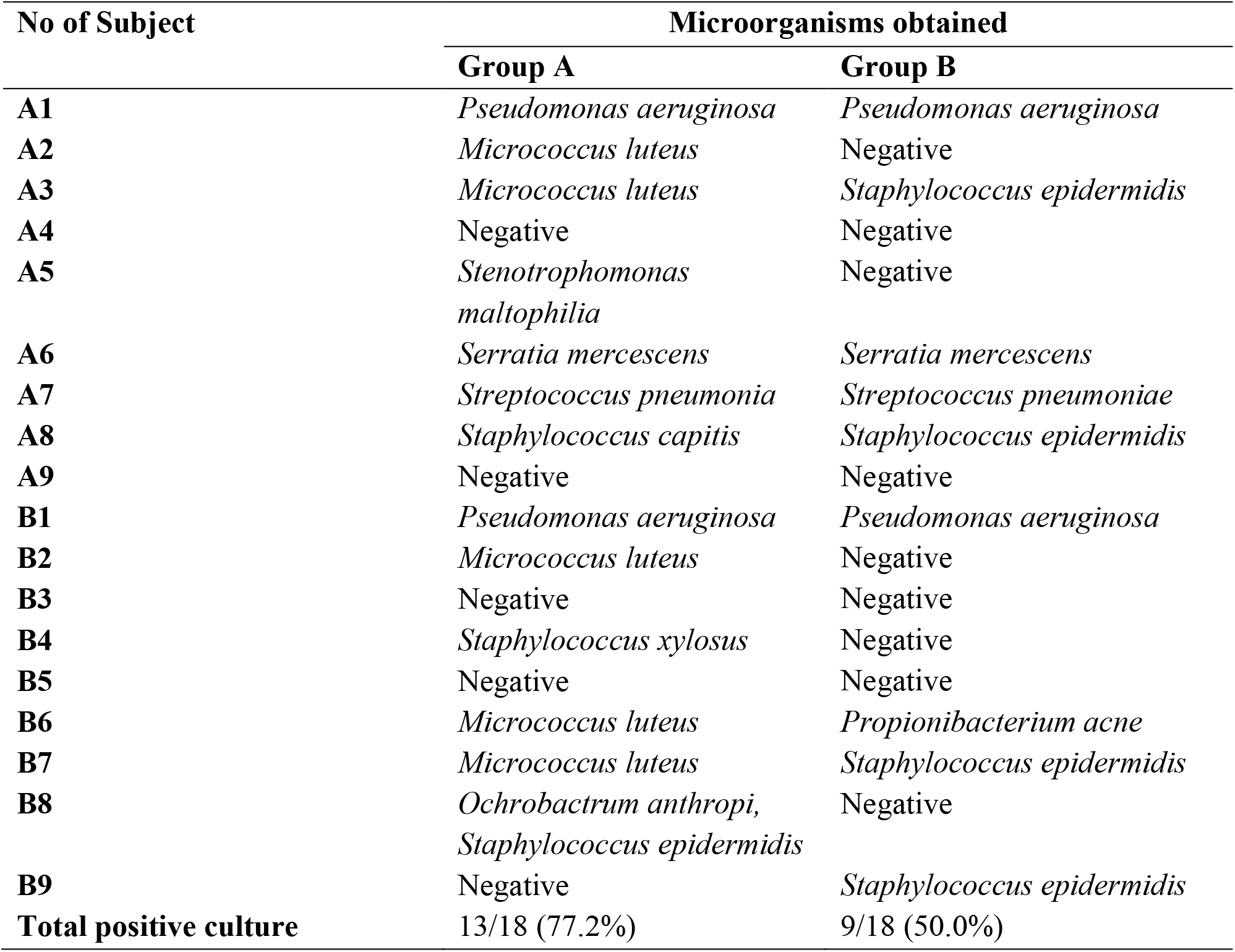
Positive Culture Using Scraping-Microhomogenization Technique and Swab Technique.

Table 3 shows the proportion of Gram according to culture in the scraping-microhomogenization technique is 6/13 (46.2%) and the proportion of Gram according to cultures in swab technique is 5/13 (38.5%). McNemar test p-value is 1.000 (p> 0.05), statistically there is no difference in the proportion of Gram and culture compatibility between the scraping-microhomogenization technique and swab technique. The Kappa suitability test value from the scraping-microhomogenization technique to the swab technique is 0.217 (fair agreement or “sufficient”). This value is lower than the expected suitability, the “good” level of compliance (Kappa 0.61-0.80). The Kappa p-value is 0.429 (>0.05) or not statistically significant.

**Table 3.** Comparison of the Proportion of Suitability of Microscopic Examination with Gram Stain and Culture on Scrapings and Smears Specimens.

|  |  | Scrapings according to culture | Scrapings are not suitable for culture | total | p* |
| --- | --- | --- | --- | --- | --- |
| Swab | According to culture | 3 (23.1%) | 2 (15.4%) | 5 (38.5%) | 1.000 |
|  | Not suitable for culture | 3 (23.1%) | 5 (38.5%) | 8 (61.5%) |  |
|  | Total | 6 (46.2%) | 7 (53.8%) | 13 (100%) |  |

\* Mc Nemar test

Table 4 shows the relationship between culture examination techniques and positive culture results. The proportion of positive cultures in the scraping-microhomogenization technique is 13/18 (72.2%), while the proportion of positive cultures in the swab technique for 9/18 (50%). McNemar test p-value is 0.219 (p> 0.05). The Kappa suitability test value from the scraping-microhomogenization technique to the swab technique is 0.333 (fair agreement or “sufficient”). This value is lower than the expected suitability, the “good” level of compliance (Kappa 0.61-0.80). The Kappa p-value is 0.114 (> 0.05) or not statistically significant.

**Table 4.** The Comparison of Positive Culture between Specimen from Scraping-Microhomogenization Technique and Swab Technique.

|  |  | Scraping-Microhomogenization |  |  | p* | Kappa | p** |
| --- | --- | --- | --- | --- | --- | --- | --- |
|  |  | Positive culture | Negative Culture | Total |  |  |  |
| Swab | Positive Culture | 8(44.4%) | 1(5.6%) | 9(50%) | 0.219 | 0.333 | 0.114 |
|  | Negative Culture | 5(27.8%) | 4(22.2%) | 9(50%) |  |  |  |
|  | Total | 13(72.2%) | 5(27.8%) | 18(100%) |  |  |  |

p\*: Mc Nemar test, p\*\*: Kappa test

## DISCUSSION

A comparative cross-sectional audit of 18 corneal scrapes undertaken at Cipto Mangunkusumo Hospital between July and November 2016 demonstrated that The proportion Gram of group A is 6/13 (46.2%), while the proportion Gram of group B is 5/13 (38.5%), which is not statistically significant (p> 0.05). The correspondence between the two techniques in terms of Gram according to culture has a Kappa suitability test value of 0.217 (fair agreement or “sufficient”) with a value of p>0.05. This can be caused by the location of a material collection which is different between the two techniques. The scraping technique takes the corneal specimen from beneath the surface and debris at the edge and base of the ulcer, while the swab technique takes the corneal specimen on the surface of the ulcer. Data on the suitability of Gram staining with low culture were also obtained from the previous descriptive studies in our hospital using corneal swabs which stated that 23.5% results of Gram-negative rods and 66.7% results of Gram-positive cocci were not in accordance with the results of their culture.^7^ One study also reported that the results of the Gram examination are incompatible with culture results in bacterial corneal ulcers, only 57% Gram staining was compatible with the culture.^12^ This can be caused by several factors such as too few specimens, the possibility of errors during specimen collections, transportation of media to the laboratory, or when the medium is examined in the laboratory. In patients who have previously been given empirical antibiotic therapy, Gram examination and culture also give negative results.^1,12^

The total number of positive cultures in this study is 14/18 (77.8%), slightly different when each technique was assessed separately (72.2% with scraping-microhomogenization technique, 50% with swab technique). The positive culture results with the scraping-microhomogenization technique in this study were similar to the positive cultures with the same technique in Diamond et al’s study, which is 71%. Additional micro-homogenization procedures theoretically can cover up the weakness of the corneal scraping technique, which is the small sample volume and uneven sample distribution among the agar culture media.^10^

Suppose the first culture result is negative in a patient with suspected bacterial infection, in that case, a repeat culture may be performed using scraping-microhomogenization, as in 33% of samples in group B. This is considered to be clinically significant and shows the superiority of the latter examination technique.^10^

The most common isolates obtained in positive culture samples in this study were bacteria Gram-positive cocci 14/22 (63%), this is in agreement with studies in India (69.4%) and Africa (68.9%). Followed by Gram-negative rods 7/22 (32%) according to studies in India (30.6%) and Africa (17%).^13^ The most bacterial species found in this study were *Staphylococcus epidermidis* (22.7%), *Micrococcus luteus* (22.7%), and *Pseudomonas aeruginosa* (18%). From a study in India, the most common bacterial species were *Streptococcus* (46.8%), *Staphylococcus* (24.7%), and *Pseudomonas* (14.9%). From a study in Africa, it was found that the most common bacterial species were *Pseudomonas* (52.5%), *Streptococcus* (20%), and *Corynobacterium* (10%).^14^

In several studies, *Staphylococcus sp* is the most common bacteria in bacterial corneal ulcers. In this study, bacteria were grown the most (22.7%). This is similar to the study in Africa, where *Staphylococcus epidermidis* was the most common type found in bacterial corneal ulcers at 27%.^13^

In one patient with risk factors for diabetes mellitus, *Ochrobactrum anthropi* bacteria were found in culture isolates using microhomogenization scrapings. Ochrobactrum anthropi belongs to the group of Gram-negative rods that have been reported as a cause of corneal ulcers and are aggressive in immunocompromised patients.^15^ This bacterium has also been reported as a cause of endophthalmitis.

There were no complications such as corneal perforation during sampling in this study. This is under the studies of Diamond et al and Krisna et al.^10,13^ Which showed that the safety level of the corneal scraping technique was quite good.

## Conclusion

The results of bacterial culture from specimens taken by the corneal scraping followed by the microhomogenization technique have a higher positive culture number (72.2%) compared to those of bacterial cultures from specimens taken with a corneal swab (50%), but were not statistically significant (p>0.05), with a Kappa value of 0.333.

The application of the corneal scraping followed by microhomogenization in specimen culture collection in patients with moderate and severe bacterial corneal ulcers can be considered because it has clinical advantages, although not statistically significant.

## Acknowledgment

The authors would like to thank Ferdy Iskandar, MD for his assitance

